# Comprehensive catalogs for microbial genes and metagenome-assembled genomes of the swine lower respiratory tract microbiome identify the relationship of microbial species with lung lesions

**DOI:** 10.1101/2023.07.25.550507

**Authors:** Jingquan Li, Fei Huang, Yunyan Zhou, Tao Huang, Xinkai Tong, Mingpeng Zhang, Jiaqi Chen, Zhou Zhang, Huipeng Du, Zifeng Liu, Meng Zhou, Yiwen Xiahou, Huashui Ai, Congying Chen, Lusheng Huang

## Abstract

Understanding the community structure and functional capacity of the lower respiratory tract microbiome is crucial for elucidating its roles in respiratory tract diseases. However, there are few studies about it owing to the difficulty in obtaining microbial samples from the lower respiratory tract. Here we collected 745 microbial samples from the porcine lower respiratory tract by harvesting 675 pigs, and constructed a gene catalog containing 10,337,194 nonredundant genes, of which only 30% could be annotated taxonomically. We obtained 397 metagenome-assembled genomes (MAGs) including 111 MAGs with high-quality. These 397 MAGs were further clustered into 292 species-level genome bins (SGBs), among which 56% SGBs are unknown with current databases. Combining with the lung lesion phenotype, we found that *Mycoplasma hyopneumoniae* strains and the adhesion-related virulence factors harboring in their genomes were significantly associated with porcine lung lesions, implying the role of adhesion and overgrowth of pathogenic *M. hyopneumoniae* in host lung diseases. This study provided important resources for the study of porcine lower respiratory tract microbiome and lung health.

## Introduction

Respiratory diseases are considered the most important health problems in pig production, which reduce pig growth rates and feed conversion efficiency, and result in significantly economic losses^1^. Diagnosis of respiratory diseases in pigs is often challenging and should rely on the combination of the evaluation of lung lesions on abattoir inspection. Respiratory disease in pigs is multifactorial and is often the result of the interplay of infectious agents and environmental factors, such as ventilation, production system and management^2, 3^. For the past few years, growing numbers of evidences have suggested that the lung microbiome has a significant impact on respiratory diseases^4, 5^. It is thus critical to reveal the community structure and function capacity of the lung microbiome for explaining its roles in maintaining lung health and in related diseases. Although the diversity and abundance of the microbiota in the lung are much lower than that in the oral and gut, culture-independent techniques such as 16S rRNA gene sequencing have demonstrated that the lung microbiota is diverse and highly individual specific^6^. And *Mycoplasma hyopneumoniae* has been identified as the etiological agent of porcine chronic respiratory (enzootic pneumonia)^7^. However, to our knowledge, the functional capacity and species-level microbial composition of pig lung microbiome remain unexplored. In addition, many unknown microbial species are yet to be characterized.

The rapid development of next-generation sequencing technologies makes it possible to infer the functional capacity of microbial communities and to explore the microbial species associated with lung diseases by shotgun metagenomic sequencing analysis. At present, the metagenomic sequencing of lung microbiota has been used to diagnose lung diseases and respiratory infections in humans. For example, Losada *et al*. used deep metagenomic sequencing to reveal more than 1000 microbial species, including bacteria, fungi, archaea, and viruses in cystic pulmonary fibrosis-induced sputum samples from patients at different ages^8^. However, a major limitation related to the utilization of metagenomic sequencing in the studies of respiratory tract microbiome is the lack of a comprehensive catalogue of reference genes/genomes for in-depth functional metagenome analysis. Although Dai *et al*. established a respiratory tract microbial gene catalogue (RMGC) containing 2.25 million non-redundant microbial genes with the microbial samples obtained from Chinese children^9^, the collection of bronchoalveolar lavage (BAL) fluid samples from participants is highly difficult in the studies of human lung microbiotas. This should not be a problem in pigs. Microbial samples can be obtained immediately from pig lungs removed from the thoracic cavity after slaughter, effectively avoiding contamination from the mouth and upper respiratory tract. However, to our knowledge, there are no previous reports concerning microbial genes or genome catalogs of the respiratory tract microbiome in pigs. This severely limits the utilization of metagenomic sequencing technology in the analysis of pig lower respiratory tract microbial community, and in the diagnosis of pig respiratory diseases. Therefore, constructing a comprehensive gene and genome catalog from a wide range of respiratory sample sources via deep metagenomic sequencing is eagerly needed for the study of pig respiratory microbiome.

In this study, we performed metagenomic sequencing on 745 microbial samples collected from the lower respiratory tract of five pig breeds (**Supplementary Table 1**). An average of 1.5 Gb clean microbial reads per sample was obtained. We constructed a comprehensive pig lower respiratory tract gene catalog (PRGC) containing 10,337,194 genes clustered at the 90% protein identity, of which 46% were unknown proteins. Using binning analysis, 397 metagenome-assembled genomes (MAGs) with medium to high-quality^10^ were obtained, which were further clustered into 292 species-level genomes. Using these comprehensive gene and genome catalogs, we characterized the diversity and functional capacities of the microbiome in the lower respiratory tract at a high resolution. Meanwhile, considering the possible correlation of the lower respiratory tract microbiome with lung diseases, we explored the association between lung microbiome and lung lesions which was considered as a trait reflecting the status of lung diseases.

## Results

### Metagenomic sequencing data of 745 microbial samples collected from the pig lower respiratory tract

We collected 745 microbial samples comprising 670 BAL samples, 74 tracheal lavage samples, and one esophageal lavage sample within 30 minutes after slaughter (see Methods) from 675 pigs composed of 618 pigs from the F_7_ generation of a mosaic population, 28 pigs from a Berkshire × Licha line, 11 Tibetan pigs, nine Erhualian pigs, and nine wild boars (**Supplementary Table 2**). These experimental pig populations included constructed experimental population (F_7_), commercial line, Chinese indigenous pig breeds (Erhualian and Tibetan) and wild boars, and were broadly representative. All DNA samples were used for shotgun metagenomic sequencing with a 150-bp paired-end strategy and generated an average of 70.30 Gbp of raw sequencing data per sample (**Supplementary Fig. 1a**). After quality control and removing host DNA contamination, an average of 1.52 Gbp clean reads per sample was obtained for subsequent analyses (**Supplementary Fig. 1b**).

Six blank samples of phosphate-buffered saline (PBS) solution that was used for lavage and six sequencing background control samples from mixed reagents were collected and used for library construction and sequencing (See Methods). Compared to tested samples, significantly low sequence numbers were obtained for these control samples (28.4∼473.2 Mb *vs.* 46.2∼149.8 Gb in raw data) (**Supplementary Fig. 2**). Furthermore, among ten bacterial species detected in control samples and for which the sum of their abundances accounted for an average of 97.9% of total microbial abundance in control samples, only three species were also identified in 745 tested samples. However, their relative abundances were < 0.9% in tested samples (**Supplementary Table 3**). Therefore, the results suggested that blanks (contaminations) are unlikely to have a substantial influence on observed microbiome profiles.

### Constructing a comprehensive gene catalog of the porcine lower respiratory tract microbiome

We implemented an assembly strategy combining both single-sample assembly and co-assembly of all sequenced samples. A total of 19,685,103 contigs with a length ≥ 500bp were obtained. On average, over 81.2% of the clean reads of each sample could be aligned to these contigs. And then, all assembled contigs were used for gene prediction. All predicted genes were first filtered with the threshold of length > 100bp. However, genes with a complete open reading frame (including start and stop codons in the Prodigal prediction) were retained even if their lengths were less than 100 bp. Furthermore, all genes were clustered to construct a non-redundant gene catalog according to their amino acid sequences at the 100%, 90%, and 50% sequence identity, resulting in the gene catalogs defined as PRGC100, PRGC90, and PRGC50. After filtered out those genes which were annotated to pig genome, the numbers of genes in the three gene catalogs described above were 13,171,977 (PRGC100), 10,337,194 (PRGC90), and 5,823,149 (PRGC50) (**Supplementary Fig. 3a**). Compared with PRGC100, the number of annotated proteins in the PRGC90 was 26% less. However, the number of annotated taxa was only decreased by 1%. Thus, the PRGC90 catalog was used for further analyses (**Supplementary Fig. 3a**).

Rarefaction curves based on the PRGC90 showed that the gene number tended to saturation when the number of samples was ≥ 200 (**Supplementary Fig. 3b**). This indicated that the sample size was enough for constructing a comprehensive gene catalog of the porcine lower respiratory tract microbiome. Unexpectedly, compared to 10,337,194 genes in the PRGC90, the number of genes at saturation was only more than six million. We inferred that this was due to the utilization of a co-assembly strategy in the assembly of contigs that acquired many genes that could not be assembled in a single sample because of low abundance, but that could be obtained in co-assembly. This phenomenon can be expected under insufficient sequencing depth^11^.

We then investigated the prevalence of predicted genes across 745 samples. The number of genes in the PRGC90 present in at least one sample was 6,554,468 (63.4%). There were 811,424 (7.8%) genes that existed in > 20% samples, and only 39,925 (0.4%) genes were present in >90% of all tested samples (**Fig. 1a**). In addition, we further analyzed the relative abundance of 6.55 million genes that existed in at least one sample, and found that most of these genes showed low-abundance, and only 1,101,865 genes had an abundance greater than the average abundance (**Supplementary Fig. 3c**). These results suggested the high diversity of microbial genes in PRGC90 and the heterogeneity of the gene composition in the swine lower respiratory tract microbiome. This should facilitate the characterization of the compositions and functional capacity of the swine lower respiratory tract microbiome regarding the PRGC90.

**Fig. 1.**
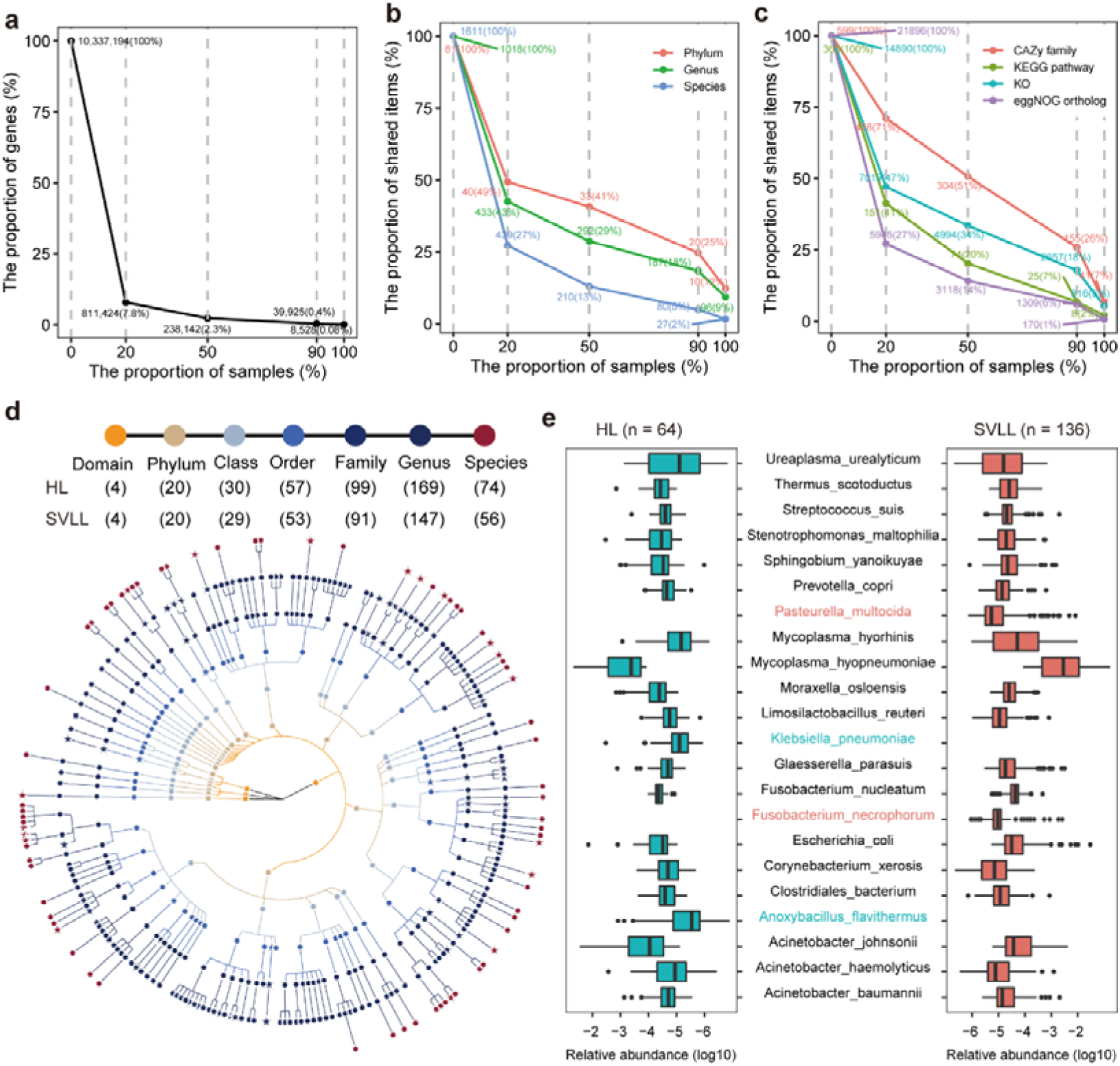
Distributions of nonredundant genes, taxa, and functional items in tested samples. (a) The numbers (percentages) of nonredundant genes in the PRGC90 shared across different proportions of 745 samples. (b) The numbers (percentages) of shared taxa among different proportions of samples at the phylum (red), genus (green), and species (blue) levels. (c) The numbers (percentages) of shared function items among different proportions of samples for CAZy family (red), KEGG pathways (olive), KEGG orthologues (cyan), and eggNOG orthologs (purple). Values next to the dots represent the numbers and proportions of items in the PRGC90 shared by each of the sample proportions 20%, 50%, 90%, and 100%. The *x*-axis indicates the proportions of samples, and the *y*-axis shows the proportions of shared items. (d) Taxonomic tree of core microbial taxa in the lung microbiome of pigs with healthy lungs (HL) and severe lung lesions (SVLL). The taxa identified in at least 95% of tested samples in each of the HL and SVLL groups were defined as core taxa. Each dot represents a taxon. The circles from the inside to outside represent the taxonomic levels from domain to species. Different colors represent different taxonomic levels. Numbers in brackets indicate the total number of core taxa at each taxonomic level. The diamond dots represent the specific core taxa in the SVLL, the star dots represent the specific core taxa in the HL, and the circle dots show the taxa shared by two groups. (e) The lung core species ranked in the top 20 in relative abundance in the HL and SVLL groups. Those top 20 species whose text colors correspond to the boxplot were specifically identified in that group. The *x*-axis shows the log10 (relative abundance) values. The box lengths indicate the interquartile range. The center line shows the median, the whiskers indicate the lowest and highest values within 1.5 times of the interquartile range from the first to third quartiles, and the points lay out outliers of the whiskers.

### Taxonomic and functional annotation of the porcine lower respiratory tract microbiome based on the PRGC90

Taxonomic annotation was performed by mapping genes in the PRGC90 to the UniProt TrEMBL database. A total of 3,286,805 (30%) genes could be annotated taxonomically. These genes were assigned to four kingdoms, 81 phyla, 1,018 genera, and 1,611 unique species. Among 1,611 microbial species, 1,484 belonged to bacteria. At the phylum level, Proteobacteria (44%), Tenericutes (31%), Firmicutes (10%), Bacteroidetes (6%), and Actinobacteria (4%) were predominant in the microbial community of the pig lower respiratory tract. A total of 10 phyla, 112 genera and 42 species were identified as virus. Among these virus taxa, *porcine parvovirus 3* and *porcine type-C oncovirus* had the highest abundances (**Supplementary Fig. 4a**). There were four phyla, 38 genera and 22 species belonging to fungi. *Pneumocystis jirovecii* were the dominant fungi species (**Supplementary Fig. 4b**). We identified nine species belonging to archaea. These nine species were classified into nine archaea genera and three phyla. Among them, *Methanobrevibacter smithii* was predominant (**Supplementary Fig. 4c**). Meanwhile, we investigated the prevalence of these annotated taxa among 745 tested samples at the phylum, genus, and species levels. Twenty phyla (25.0% of all phyla identified) were detected in more than 90% of the tested samples (**Fig. 1b**). In contrast, only 187 genera (18.0%), and 80 species (5.0%) were detected in > 90.0% of tested samples, suggesting the heterogeneity of microbial composition in the swine lower respiratory tract. It was worth noting that most of the microbial taxa (>97% species) with high prevalence belonged to bacteria. Virus, fungi and archaea species only accounted for 0.2% of total abundance (**Supplementary Fig. 4d**), so we focused on bacterial taxa in the subsequent analyses. Genes in the PRGC90 were annotated to Carbohydrate-Active enzymes (CAZymes), Kyoto Encyclopedia of Genes and Genomes (KEGG), and eggNOG to obtain functional capacity information. A total of 164,626 genes were annotated to 599 unique CAZy families, 2,697,548 genes to 365 KEGG pathways, and 568,498 genes to 21,986 eggNOG orthologs (**Supplementary Table 4**). Similar to the prevalence of microbial taxa, 26.0% CAZy families, 18.0% KOs, 7.0% KEGG pathways, and 6.0% eggNOG orthologs were present in more than 90.0% of all the tested samples (**Fig. 1c**). The low percentages of shared functional capacities indicated the heterogeneity of functional capacities of the swine lower respiratory microbiome.

The composition and function capacities of the lung microbial community are influenced by environmental factors^12^. We then explored the distribution of core species of the lung microbial community in the five pig populations used in this study. The microbial species detected in > 95% of tested samples were defined as the core species of pig lung microbiome. A total of 68, 72, 73, 76, and 118 core species were identified in the F_7_, Berkshire × Licha line, Tibetan pigs, Erhualian pigs, and wild boars, respectively. We focused on the core species whose abundances were ranked in the top 20 in each population. Eight species were identified in the top 20 lists in all five populations, including *M. hyopneumoniae*, *Fusobacterium nucleatum*, *Moraxella osloensis*, *Acinetobacter johnsonii*, *Prevotella copri*, *Escherichia coli*, *Streptococcus suis*, and *Glaesserella parasuis* (**Supplementary Fig. 5**). Chinese indigenous pig breeds (Erhualian and Tibetan) and wild boars had more population-specific core species (**Supplementary Fig. 5**). There were eight, six, six, and two species that were specifically identified in the top 20 lists of wild boars, Erhualian pigs, Tibetan pigs, and Berkshire × Licha pigs, respectively.

There were 69 pigs from which both tracheal and BAL samples were collected. This greatly facilitated the comparison of the microbial compositions between tracheal and BAL samples. We found that the microbial composition of tracheal samples tended to show the higher α-diversity in the index of observed species although the difference in Shannon index did not achieve significance level (**Supplementary Fig.6a**). Significant difference was also observed in the β-diversity based on the Bray-curtis distance and the principal coordinate analysis (PCoA). The microbial composition of tracheal samples had a significantly higher β-diversity compared to that of BAL samples (*P* = 0.0018) (**Supplementary Fig.6b**). Bacterial species also showed different abundances between tracheal and BAL samples. There were five species whose relative abundance was listed in the top 20 for each of tracheal and BAL sample types. (**Supplementary Fig. 6c**).

BAL samples used in this study were collected from 64 healthy pigs (HL, lung- lesion score < 0.75) and 606 pigs with lung lesions. We compared the distribution of core taxa between HL pigs (n = 64) and pigs with severe lung lesions (SVLL, lung- lesion score >3) (n = 136). As shown in Fig. 1d, 74 core microbial species were identified in HL pigs. SVLL pigs shared a large portion of common core taxa with HL pig, but had a lower number of core species (56 species) (**Fig. 1d**). Among the top 20 species in relative abundance, some pathogenic microorganisms were identified in both HL and SVLL pigs, including *M. hyopneumoniae*, *Mycoplasma hyorhinis*, *G. parasuis*, *S. suis*, and *Acinetobacter baumannii* (**Fig. 1e**), but the relative abundances of these pathogens were higher in pigs with severe lung lesions (See the details below). Additionally, we analyzed core KEGG pathways (existing in > 95% of the samples) in HL and SVLL pigs. We identified 18 core KEGG pathways in HL pigs and 19 core KEGG pathways in SVLL pigs. Among them, sixteen core KEGG pathways were detected in both pig groups, implying that core KEGG pathways were similar across HL and SVLL pigs (**Supplementary Fig. 7**). We also found that the core KEGG pathways specifically identified in the SVLL group were related to microbial pathogenesis, including bacterial motility proteins and prokaryotic defense system.

### Constructing metagenome-assembled genomes from metagenomic sequencing data of swine lower respiratory tract microbiome

Due to the high proportion of host DNA contamination in the lower respiratory tract microbial samples^13^, few studies have been able to reconstruct microbial genomes from metagenomic sequencing data of the lower respiratory tract microbiome. In this study, 2,056 MAGs with completeness ≥ 50% and contamination ≤ 10% were recovered by a metagenomic binning procedure using contigs generated by both single-sample assembly and co-assembly. The 2,056 MAGs were clustered into non- redundant 397 MAGs based on 99% average nucleotide identity (ANI) (**Supplementary Table 5**). Among these 397 MAGs, 286 MAGs showed medium- quality (at least 50% completeness and less than 10% contamination), including 218 MAGs having a quality score (QS) ≥ 50 (completeness - 5*contamination)^10^ and two MAGs having completeness ≥ 90%, but contamination >5.0%. The other 111 MAGs were near-complete (completeness ≥ 90% and contamination ≤ 5.0 %) (**Supplementary Fig. 8a** and **8b**), among which 20 MAGs had the 5S, 16S, 23S rRNA genes, and at least 18 tRNAs that complied with the MIMAG standards for the ‘high quality’ MAG set by the Genomic Standards Consortium^14^. Prevalence analysis found that most of MAGs showed low prevalence in 745 metagenomic sequencing samples. Only eight MAGs were identified in >90% of tested samples, and 170 MAGs were present in at least 20% of tested samples, further suggesting that the microbial composition of the swine lower respiratory tract was highly heterogeneous across samples, and more samples would benefit the reconstruction of microbial genomes. Certainly, other reasons, such as different sequencing depth across samples and the complications of the assembly process of MAG re-constructions could not be excluded.

All 397 non-redundant MAGs were subjected to taxonomic annotation using GTDB-Tk^15^. Among them, 395 MAGs could be classified as bacteria (16 phyla), and the other two MAGs were annotated to *ISO4-G1* (Thermoplasmatota) and *Methanobrevibacter A sp900769095* (Methanobacteriota) belonging to methanogens archaea. However, only 46% (186) of MAGs could be assigned to 131 unique species (**Fig. 2a** and **2b**). According to the taxonomic annotation, the largest number of MAGs (n = 122) was assigned to Proteobacteria, followed by Bacteroidetes (n = 100) and Firmicutes (n = 91). To determine the phylogenetic relationships of these MAGs, a maximum likelihood phylogenetic tree of 397 MAGs was constructed in PhyloPhlAn based on 400 universal marker genes (**Fig. 2c**). MAGs belonging to Proteobacteria showed a wide distribution in the phylogenetic tree, indicating high phylogenetic diversity.

**Fig. 2.**
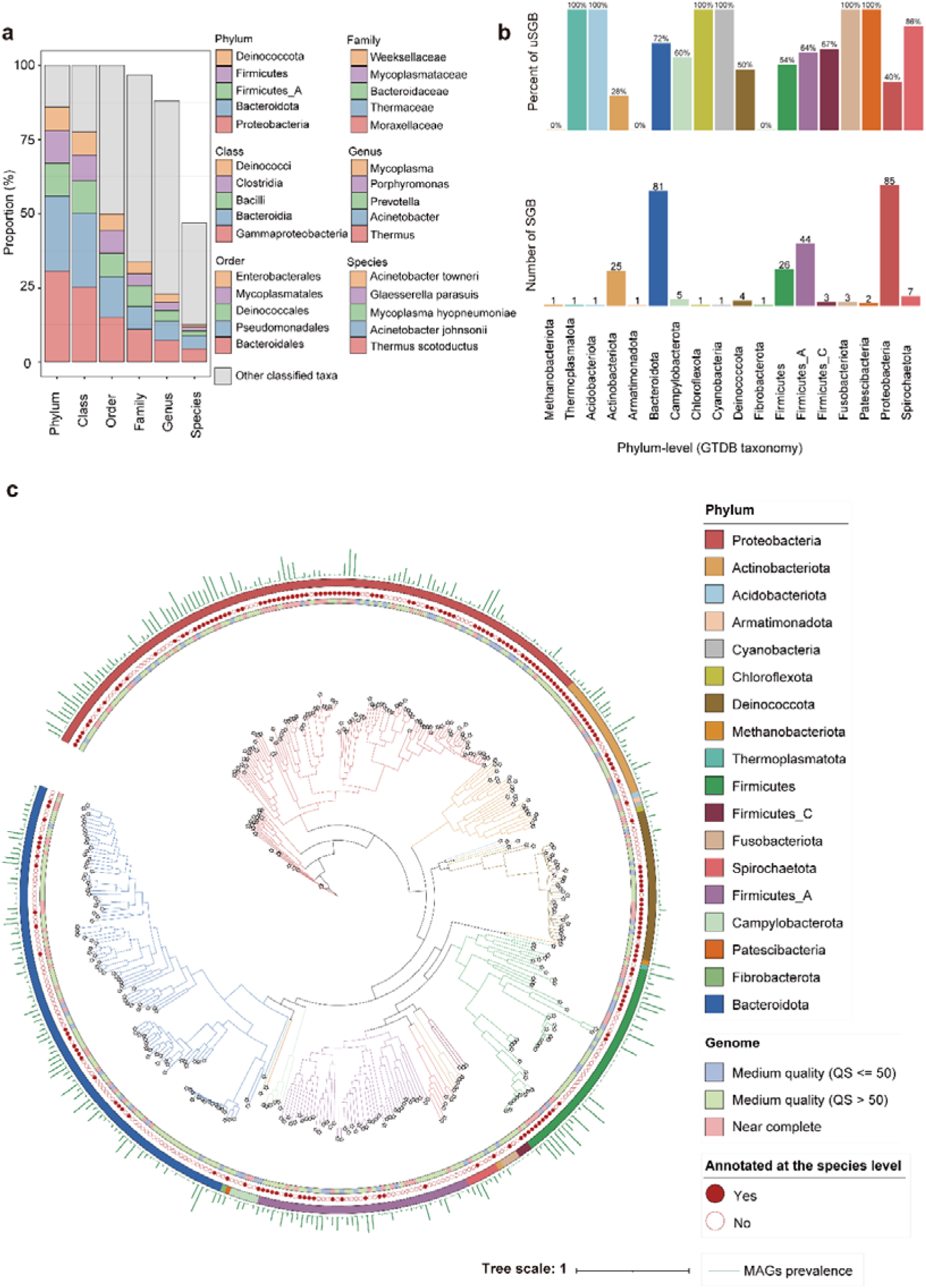
Taxonomic annotation and phylogeny of 397 metagenome-assembled genomes (MAGs) (a) Taxonomic composition of 397 MAGs. The colored boxes indicate the proportion of MAGs annotated to each taxon. Only five taxa with the highest proportions of MAGs at each level are shown, and other taxa were classified into ‘other classified taxa’. (b) The number of species-level genome bins (SGBs) in each phylum and the percentage of unknown SGBs (uSGBs) in each phylum. The SGBs that could not match a reference genome using GTDB-tk were defined as uSGBs. (c) Phylogenetic tree of 397 MAGs constructed in swine lower respiratory tract microbiome. From the inner to outer circles, the first circle indicates the quality of MAGs (medium quality (QS ≤ 50), medium quality (QS > 50) or near-complete), the second circle shows the MAGs that could be annotated to species or not, the third circle indicates the corresponding phyla that MAGs were belonged to, and the green bars in the fourth layer represent the prevalence of MAGs in 745 tested samples.

All 397 MAGs were further clustered into 292 species-level genome bins (SGBs) at the threshold of 95% ANI. Among these 292 SGBs, 56% (n = 163) did not match any available genomes within GTDB-Tk, and therefore, were defined as unknown SGBs (uSGBs) (**Fig. 2b**). Just like the condition for MAGs, the largest number of SGBs belonged to Proteobacteria (29%, n = 85), followed by Bacteroidota (28%, n = 81) and Firmicutes_A (15%, n = 44). Among the top three SGBs with the largest number of MAGs, the SGBs annotated to *T. scotoductus* and *A. johnsonii* contained 18 and 17 MAGs, followed by the SGB belonging to *M. hyopneumoniae* that was comprised of seven MAGs. All 163 uSGBs could be assigned to a known order, and 83% were annotated at the genus level. This suggested that a substantial portion of uSGBs might be novel bacterial species.

### Pan-genome and phylogenetic structure of *M. hyopneumoniae* genomes

Among 2,056 MAGs (before dereplication) constructed in this study, the largest number of MAGs (n = 262) was obtained for *M. hyopneumoniae*. As one of main pathogens for lung diseases^7^, *M. hyopneumoniae* is not easy to culture because it grows slowly compared to other bacteria^16^. Culture-independent approaches, such as high-throughput metagenomic sequencing have been demonstrated to be an effective method for acquiring more bacterial genomes by metagenomic binning. Here we obtained a total of 285 *M. hyopneumoniae* genomes that were used for pan-genome analysis, including 262 MAGs constructed in this study with high quality (see Methods), and 23 *M. hyopneumoniae* genomes downloaded from the NCBI RefSeq database (**Supplementary Table 6**). A total of 3,121 pan-genes were obtained from these 285 *M. hyopneumoniae* genome via the Roary pipeline. Furthermore, 401 genes present in > 95% of the genomes were defined as core genes. In addition, 2,361 (76%) genes were only detected in less than 42 (15.0%) genomes, indicating the strain-specific gene composition. The number of pan-genes in *M. hyopneumoniae* genomes did not tend to saturation, suggesting an increasing repertoire of gene content within population (**Fig. 3a**). The reconstructed MAGs from each sample showed a much greater size of the collective pan-genes than genomes downloaded from the database. This result suggested that *M. hyopneumoniae* genomes reconstructed in this research have greatly expanded the genome diversity of *M. hyopneumoniae*. We next examined functional categories of both core genes and pan-genes. Based on the annotations of cluster of orthologous groups (COGs) and KEGG pathways, there were no significant differences in functional categories between pan-genes and core genes (**Fig. 3b** and **Supplementary Fig. 9a**). There were 2,539 (81%) pan-genes that could be annotated to COGs. Among them, 66% of COGs were classified into 20 COG categories, and the other 34% of COGs (n = 856) could not hit a known COG category (**Fig. 3b**). In contrast, 395 (99%) core genes had a match to eggNOG orthologs, and belonged to 17 COG categories. KEGG pathway analysis found multiple pathways related to microbial virulence in both pan-genes and core genes, including bacterial secretion system, protein export, and quorum sensing pathways (**Supplementary Fig. 9a**). This suggested that the virulence-related functions were important parts of *M. pneumoniae* functional capacities.

**Fig. 3.**
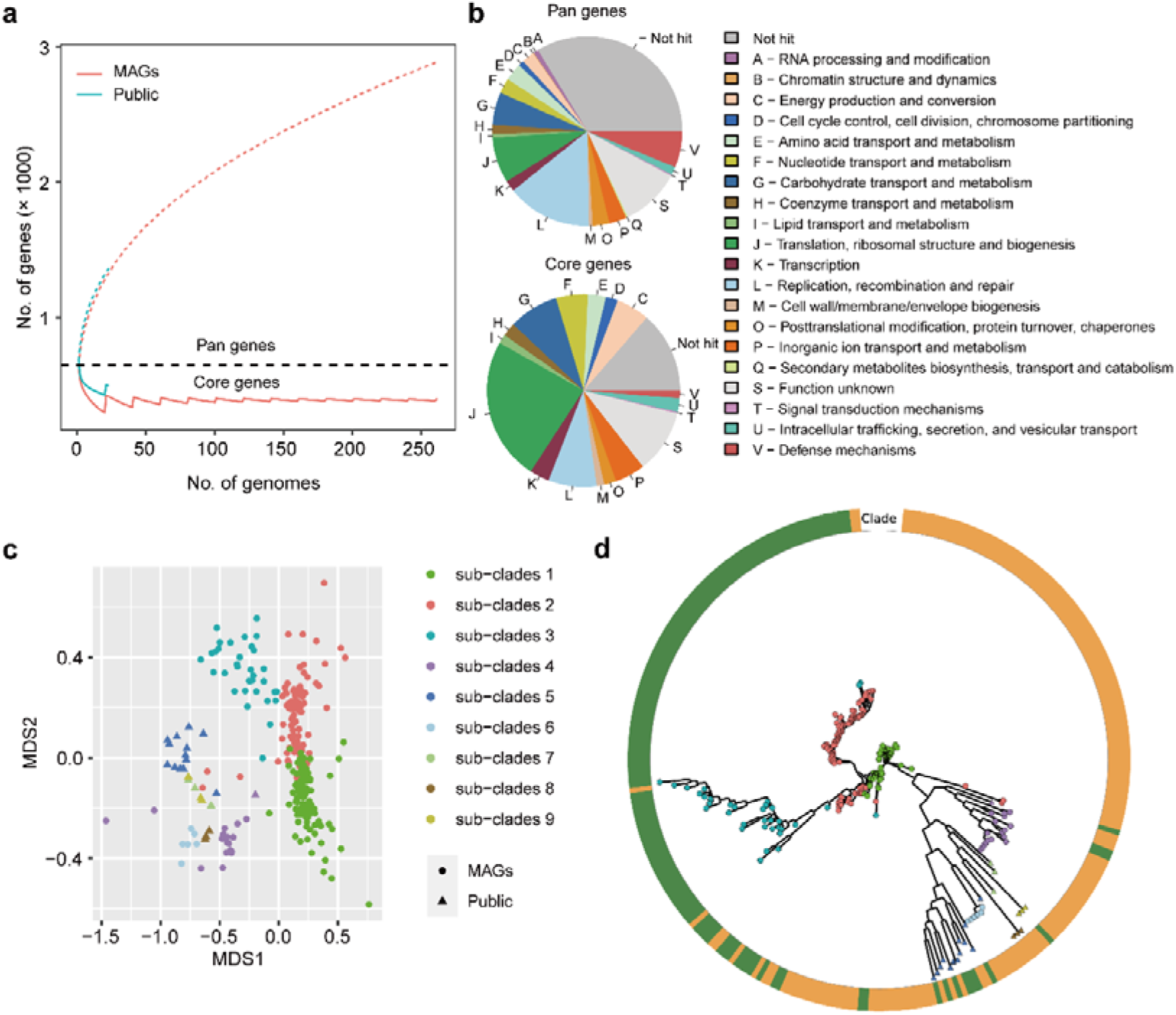
Pan-genome analysis of 285 *M. hyopneumoniae* genomes. (a) Gene accumulation curves for pan-genes and core genes in 262 MAGs reconstructed in this study, and 23 *M. hyopneumoniae* genomes download from the NCBI RefSeq database. (b) The clusters of orthologous groups (COG) of pan-genes and core genes. Not hit represents the genes could not be aligned to any COG category in the database. (c) Classical multidimensional scaling (MDS) analysis for 285 *M. hyopneumoniae* genomes based on average nucleotide identity (ANI) distance matrix. Each dot represents a MAG, and triangles represents publicly available reference genomes of *M. hyopneumoniae*. Different colors of dots indicate genetic sub-clades (sub-clades 1-9) inferred by rhierBAPs. (d) Maximum likelihood phylogenetic tree of all 285 *M. hyopneumoniae* genomes. The outer ring is colored according to the clades that were defined based on the partitioning around medoid (PAM) algorithm (green and orange represent Clade L and Clade L, respectively). The colors and shapes of dots in the tree are consistent with that in (c).

The phylogenetic structure of *M. hyopneumoniae* genomes was characterized by applying a phylogenetic analysis to 285 genomes. Based on the genomic ANI distances using the partitioning around medoids (PAM) algorithm, 285 genomes could be clustered into two clades (predicted strength exceeded 0.8 for k = 2) (see Methods). These two clades might belong to two sub-species of *M. hyopneumoniae* at the taxonomic level. We further explored the diversity of *M. hyopneumoniae* genome sub- clades. Nine sub-clades were obtained by Bayesian clustering based on multiple sequence alignments of 401 core genes. The number of *M. hyopneumoniae* genomes included in each sub-clade ranged from three (for sub-clade 9) to 111 (for sub-clade 1). All 23 *M. hyopneumoniae* genomes downloaded from the NCBI RefSeq database were present in five out of nine sub-clades, and all showed higher genetic similarity (closer NMD distance) than MAGs reconstructed in this study (**Fig. 3c**). There were five sub-clades containing the retrieved MAGs and exhibiting high genetic distance. A maximum-likelihood phylogenetic tree was constructed based on multiple sequence alignments generated from 401 core genes, and revealed that the genomes from the public database spanned a limited phylogenetic space, whereas MAGs exhibited a broad phylogenetic structure (**Fig. 3d**). Therefore, the reconstructed MAGs expanded the genetic diversity of *M. hyopneumoniae* genomes greatly, and could facilitate further investigation of the biological functions of *M. hyopneumoniae* at the strain level.

Next, we investigated whether the functional repertoire was differed between the two sub-species of *M. hyopneumoniae* by performing functional annotation. We found that the strains belonging to clade L harbored significantly more eggNOG and KEGG orthologs, while the number of virulence factor genes (VFGs) was not significantly different between two sub-species (**Supplementary Fig. 9b**). Furthermore, clade L exhibited a much higher density of functional genes. All these results demonstrated the distinct functional repertoires of two *M. hyopneumoniae* sub-species.

### Virulence factor genes and their host bacteria

Virulence factors (VFs) refer to gene products that enable microorganisms to survive on or within a given host and thereby enhance their disease-causing potentials^17^. In the PRGC90, we identified 825,543 open reading frames (ORFs) encoding virulence factors by blasting against the virulence factors database (VFDB)^18^. These ORFs could be classified into 17,278 VFGs belonging to 1,180 VF types (e.g., P97 and Hemolysin) and 14 functional VF categories (e.g., adherence and exotoxin). The main functional categories of these 17,278 virulence factor genes (VFGs) included immune modulation (22.2% of the total number), effector delivery system (15.9%), nutritional/metabolic factor (15.5%), adherence (13.0%), and motility (12.2%) (**Supplementary Table 7**). We further aligned the sequences of VFGs to the UniProt TrEMBL database to identify their potential host bacteria. At the genus level, 17,278 VFGs were widely distributed in 563 bacterial genera belonging to 30 phyla, especially *Pseudomonas* and *Acinetobacter*. At the species level, a total of 670 host bacterial species were identified (**Supplementary Table 8**). The distribution of host species in which the numbers of VFGs they harbored were ranked in the top 100 are shown in **Fig. 4**. *E. coli* carried the largest number of 3,026 VFGs.

**Fig. 4.**
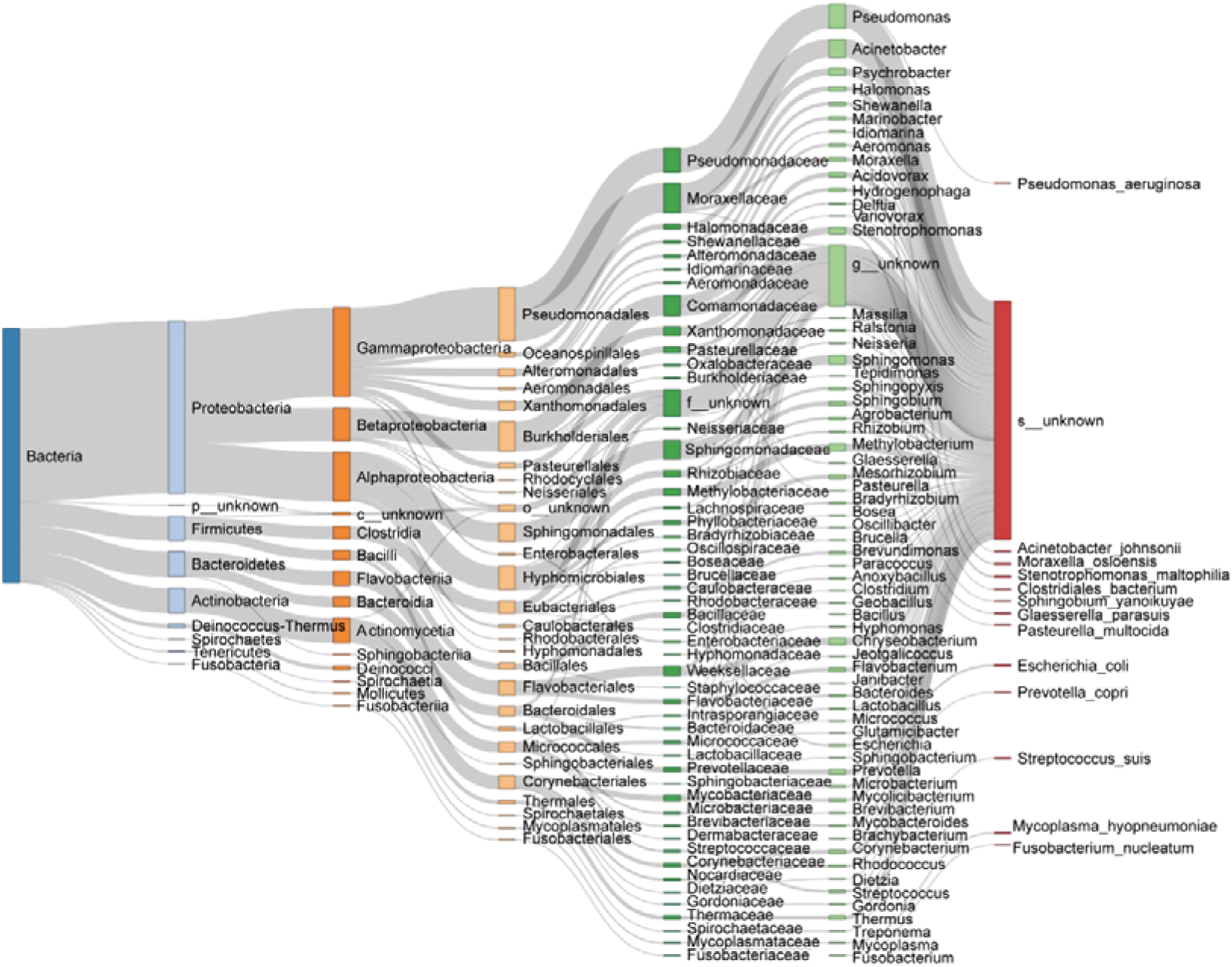
Host bacteria of virulence factor genes (VFGs) The distribution of host bacteria in which the numbers of VFGs harbored were ranked in the top 100. The colored rectangles from left to right represent different taxonomy levels from domain to species. The widths of the rectangles indicate the number of VFGs.

We performed an in-depth analysis of the distribution about VFGs in MAGs. A total of 11,099 VFGs were identified in 397 MAGs. The highest number of VFGs (995) was detected in the MAG_290 that was annotated to *Mesorhizobium*, while the MAG_366 annotated to *M. hyopneumonia* harbored the least number of 19 VFGs. As *M. hyopneumoniae* has been known as a primary pathogenic microorganism that can cause swine enzootic pneumonia by colonizing epithelial cilia^19^, we deeply investigated the distribution of VFGs in 285 *M. hyopneumoniae* genomes mentioned above. A total of 229 VFGs were identified in these genomes. However, we did not observe any discrepancies in the distribution of VFGs between two sub-species (**Supplementary Fig. 9c**).

### Identifying bacterial taxa and potential function capacities associated with lung lesions

To minimize the influence of various environmental factors on experimental pigs, F_7_ pigs of the mosaic population used in this study were raised in the same farm with uniform conditions including the same commercial feed. This provided an opportunity to better understand the relationship of the lung microbiome with lung lesions using the PRGC90 and MAG catalogs. BAL samples from 613 F_7_ pigs of the mosaic population were used in the analysis and were divided into four groups according to the lung lesion scores (**Fig. 5**), including healthy (HL) (n = 51), slight lung lesion (SLL) (n = 217), moderate lung lesion (MLL) (n = 218), and severe lung lesion (SVLL) (n = 127) groups (see Methods). We first compared the α- and β-diversity of the lung microbiota across the four groups. The HL group exhibited the highest α- diversity of lung microbiota based on the indices of Shannon and observed species, and the α-diversity decreased following the increased severity of lung lesions (**Fig. 6a**). The β-diversity of the lung microbial community among groups was also significantly different, and the similarity of the lung microbial compositions among individuals in the HL group was higher than that in other groups (**Fig. 6b**). Thus, the changed diversity of lung microbial compositions among groups suggested an association of lung microbiota with lung lesions.

**Fig. 5.**
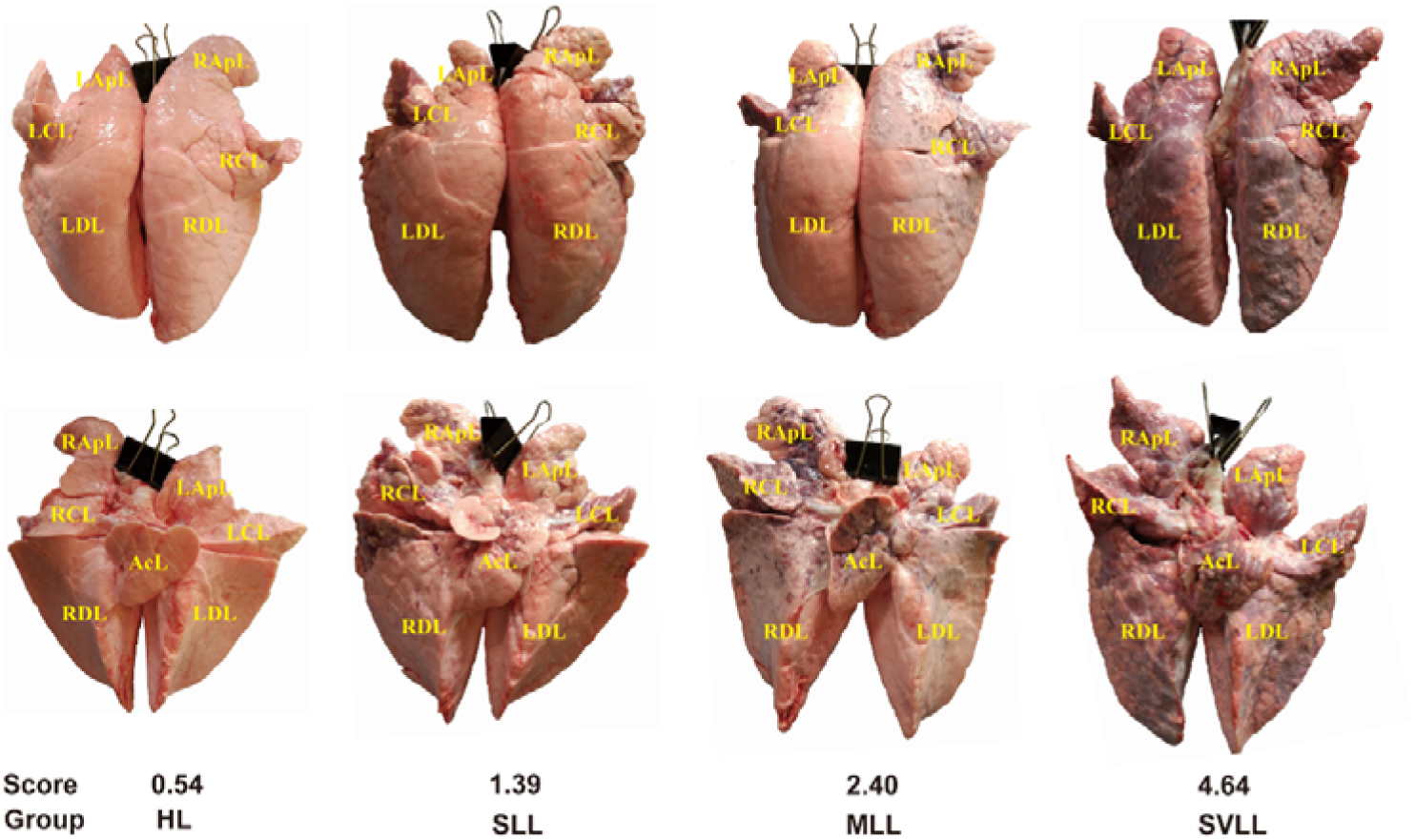
The scoring of lung lesions for grouping of 613 F_7_ pigs. The anterior and posterior views of lung were photographed for each pig using a digital camera. Six sections of the anterior lung including left apical lobe (LApL), right apical lobe (RApL), left cardiac lobe (LCL), right cardiac lobe (RCL), left diaphragmatic lobe (LDL), and right diaphragmatic lobe (RDL), and seven sections of the posterior including LApL, RApL, LCL, RCL, LDL, RDL, and accessory lobe (AcL) were scored using the methods described by (Zhang et al., 2019, and Methods). Pigs with a final score of lung lesions ranging from 0 to 0.75 were classified into the healthy lung group (HL, n = 51), from 0.75 to 1.50 into slight lung-lesion group (SLL, n = 217), from 1.5 and 3.00 into the moderate lung-lesion group (MLL, n = 218), and > 3 into the severe lung-lesion group (SVLL, n = 127).

**Fig. 6.**
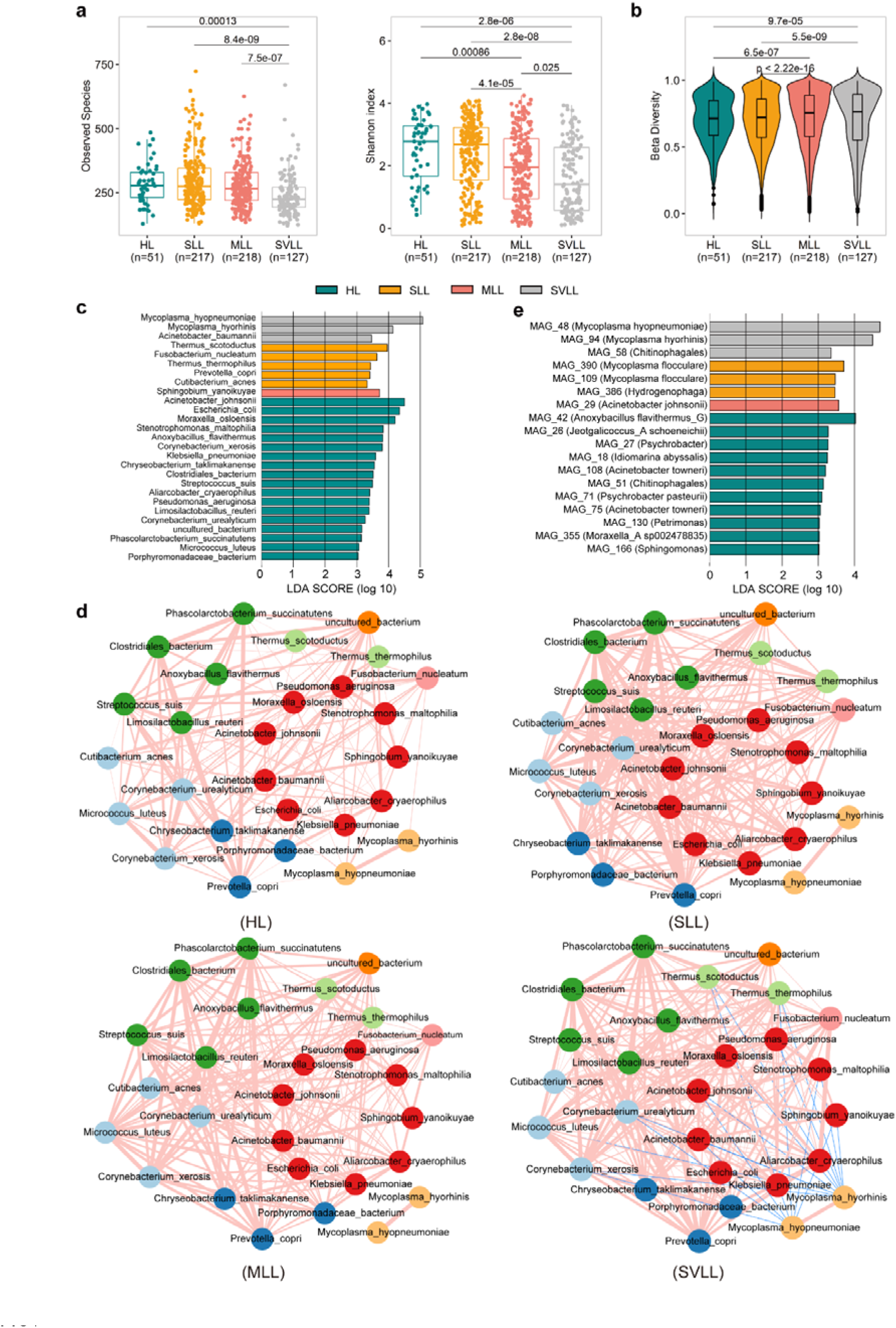
Association of the lung microbial composition with swine lung lesions. (a) Comparison of the α-diversity of the lung microbiome (observed species and Shannon index) among four pig groups with different lung lesions in the F7 population. Healthy lungs (HL, dark green), slight lung-lesion (SLL, orange), moderate lung-lesion (MLL, light red), and severe lung lesions (SVLL, light grey). Wilcoxon tests were used for the comparison of the α-diversity values across different groups. The legend for the box plots is as same as that for Fig. 1e. (b) Comparison of the β-diversity of the lung microbial composition among four pig groups with different lung lesions based on pair-wise Bray–Curtis distances. Wilcoxon tests were used for the comparison of the microbial composition across the four groups. The legend for box plots is the same as in Fig. 1e. (c) The core species enriched in each pig group by linear discriminant analysis effect size (LEfSe) analysis. (d) Co- occurrence network of 27 differential bacteria species. Interactions between species (nodes) are represented by connecting lines (edges), and each node is coloured according to the phylum it belongs to. The colors of the edge lines represent positive (light red) or negative (dark blue) interactions, and the width of edge represents the magnitude of the Spearman correlation coefficient between species. (e) The MAGs enriched in each pig group by LEfSe analysis. The corresponding taxonomic annotations of MAGs are shown in the brackets using GTDB-tk.

We then identified the microbial species associated with lung lesions. To avoid the effect of low prevalence on the statistical analysis, we focused on the core microbial species. We observed a broad dissimilarity of microbial signatures with 27 core bacterial species showing significantly different abundances among four pig groups with different severity of lung lesions (Fig. 6c). Among these differential species, the relative abundances of two *Mycoplasma* species clearly increased following the severity of lung lesions (**Supplementary Fig. 10a**) and had the most significant association with pig lung lesions (**Fig. 6c**). We further constructed a co- occurrence network to determine the relationships among 27 differential species described above (**Fig. 6d**). Overall, the co-occurrence network could be obviously divided into two sub-modules. One module only contained two *Mycoplasma* species and showed significantly less connections with the other 25 species in the HL, SLL and MLL groups. The other module contained 25 species that were positively connected each other. We observed two significantly different signatures of the interaction network among four pig groups: 1) the connections among 27 bacterial species in the health group (HL) were significantly less than in the other three groups (the average number of edges per node: 10.0 *vs.* 16.0, *P* < 0.01), suggesting fewer interactions among bacterial species in healthy pigs. 2) Significantly negative interactions were observed between *Mycoplasma spp.* and 11 differential species in the other module in pigs with severe lung lesions (the SVLL group). However, in the HL, SLL and MLL groups, few interactions (only several positive connections) were identified between *Mycoplasma spp.* and the species in the other module (**Fig. 6d**). Considering the most significant association with lung lesions and the increased abundance with the severity of lung lesions (**Fig. 6c** and **Supplementary Fig. 10a**), we speculated that the overgrowth (high abundance) of *Mycoplasma spp.* should influence the abundance of other non-*Mycoplasma* species, and the changed microbial ecosystem was associated with pig lung lesions.

The construction of MAGs in the swine lower respiratory tract microbiome allowed us to identify bacterial strains related to lung lesions. A total of 18 MAGs showed different enrichments among four pig groups, including 11 MAGs enriched in healthy pigs. Three MAGs were significantly enriched in the SVLL group, two of which were annotated to *Mycoplasma* (*M. hyopneumoniae* and *M. hyorhinis*). There were also two *M. flocculare* MAGs having higher abundances in SLL pigs (**Fig. 6e**). This further suggested that bacterial species from *Mycoplasma* were crucial to pig lung lesions. Interestingly, seven out of 397 MAGs constructed in this study belonged to *M. hyopneumoniae*. However, only MAG_48 showed the significant enrichment in pigs with severe lung lesions. This indicated that not all *M. hyopneumoniae* strains were associated with lung lesions. We found that, compared to the other six *M. hyopneumoniae* MAGs, the MAG_48 had the highest abundance in the lower respiratory tract microbiome (**Supplementary Table 5**), suggesting that the high abundance of the MAG_48 strain of *M. hyopneumoniae* was associated with the lung lesions.

We further investigated the changes in functional capacities of the lower respiratory tract microbiome following the severity of lung lesions, especially focusing on VFGs and KEGG pathways. The linear discriminant analysis identified 28 VF types showing different abundances among the four pig groups (**Supplementary Fig. 10b**). Thirteen VF types were significantly enriched in the SVLL group. Eight out of these 13 VF types were derived from *M. hyopneumoniae*. Among these eight VF types, P146, P102, P65, P159, P216, and P97/P102 paralog family are known as adhesins, which should promote the adhesion of *M. hyopneumoniae* to the host lung cells (**Supplementary Fig. 10b**). Importantly, MAG_48 carried all these six VF types. There were also 10 VF types significantly enriched in the HL group. The host bacterial species of these 10 VF types were negatively correlated with two *Mycoplasma spp.* in the SVLL group (**Fig. 6d**), including *S. yanoikuyae* harboring PspA, *M. osloensis* harboring MymA operon, and *S. maltophilia* harboring Type IV pili (**Supplementary Table 9**). This suggested that the enrichments of VF types in the HL group may be due to the increased abundances of their host bacterial species. Thirty-four KEGG pathways showed significantly different enrichments between healthy and lung lesion pigs (**Supplementary Fig. 10b**), highlighting the changes in functional capacities of the pulmonary microbiome in lung lesions. Three out of four functional pathways significantly enriched in the SVLL group were related to cell communications, including proteoglycans, exosome, and CD molecules. Specially, CD molecules are related to cell adhesion^20^, and proteoglycans can regulate cellular processes, such as adhesion^21^. This result suggested the role of bacterial adhesion (*M. hyopneumoniae* adhesion as described above) in lung lesions.

To further confirm the role of bacterial strains in lung lesions across pig populations, we compared the abundances of 397 MAGs in the lung microbiome between healthy pigs (n = 11) and lung lesion pigs (lung-lesion scores >0.75, n = 17) in the Berkshire × Licha cross line (Methods). Interestingly, six MAGs belonging to *M. hyopneumoniae* were significantly enriched in pigs with lung lesions with the significance values (LDA) ranked in the top six (**Supplementary Fig. 11a**), including the MAG_48 which was also most significantly enriched in pigs with severe lung lesions in the F_7_ population. Furthermore, another *M. hyopneumoniae* strain (MAG_5) with the highest abundance in the Berkshire × Licha cross line showed the most significant association with lung lesions. All these results confirmed the significant association of *M. hyopneumoniae* strains with pig lung lesions. The VF types enriched in the lung microbiome of lung lesion pigs were also similar to that obtained in the F_7_ population (**Supplementary Fig. 11b**). P97/P102 paralog family, P102, P65, P146, P159, P216, and P97 known as adhesins were significantly enriched in lung lesion pigs. More importantly, six lung lesions-associated *M. hyopneumoniae* MAGs carried at least five of these seven adhesin VF types (**Supplementary Table 10)**. This result provided an important clue that the adhesion of *M. hyopneumoniae* strains to host lung cells, which might be mediated by virulence factors, may play a vital role in lung lesions.

Overall, we concluded that adhesion-related virulence factors carried by *M. hyopneumoniae* strains might promote the adhesion of *M. hyopneumoniae*. The overgrowth of *M. hyopneumoniae* strains (high abundances) decreased the relative abundances of other bacterial species and the diversity of the lower respiratory tract microbial community, and was finally associated with the lung lesions of pigs.

## Discussion

Here we conducted a large-scale metagenomic sequencing of microbial samples from the swine lower respiratory tract and constructed microbial gene and MAG catalogs of the swine respiratory tract microbiome. As an application of these catalogs, we found a significant correlation of *M. hyopneumoniae* strain with porcine lung lesions. We also investigated the population genomic structure of *M. hyopneumoniae* and found that *M. hyopneumoniae* exhibited a high genetic diversity in the swine respiratory microbiome. To our knowledge, these are the first microbial gene and MAG catalogs of the respiratory microbiome in pigs. It provides important genomic resources for the studies about the respiratory microbiome and has yielded new insights into the relationships between the lung microbiome and lung lesions.

To our knowledge, only Dai *et al*.^9^ constructed a respiratory microbial gene catalog containing 2.3 million genes in 247 children (both healthy children and the children infected with *Mycoplasma pneumonia*). Notably, there are few studies about metagenomic sequencing of the lower respiratory tract microbiota^22^. This is primarily due to the following reasons: 1) It is difficult to collect microbial samples from the lower respiratory tract due to its complicated structure^23^. There is a high risk of contamination from the upper respiratory tract microbiota^24^. 2) Given the low biomass of the lower respiratory tract microbiota, it is difficult to obtain sufficient DNA for next-generation sequencing^25^. More importantly, a high proportion (over 90%) of host DNA contamination has existed. This results in a small amount of valid microbial sequencing data^13^. In this study, tracheal lavage and BAL samples were obtained by separating lungs and trachea from pig thoracic cavity after slaughter. This avoided the contamination by microbiota from the upper respiratory tract. An unprecedented number of more than 10.3 million of microbial genes which exhibited the high diversity and representation of the PRGC90 catalog, which was achieved by greatly increasing the sequencing depth to an average of 70.3 Gb per sample (**Supplementary Fig. 1a**) and using a broad sample source.

Several researches have investigated the lung microbial community in pigs^26–28^. However, most of these studies were performed based on the 16S rRNA gene sequencing analysis that could only describe the microbial composition at the taxonomy level. Consistent with our previous report using 16S rRNA gene sequencing^26^, *Prevotella, Streptococcus* and *Mycoplasma* were the dominant genera. However, *Methylotenera*, *Phyllobacterium*, and *Lactobacillus* showing high abundances in our previous study were not predominant in this study. This should mainly attribute to different pig populations and sequencing methods. However, consistent with the results of this study, Siqueira *et al.* also found that *M. hyopneumoniae*, *M. hyorhinis*, *S. suis*, *G. parasuis*, and *E. coli* had high abundances in the lung microbiome of healthy pigs using shotgun metagenomic sequencing^27^.

More than 17,000 VFGs identified in the PRGC90 catalog should improve our understanding of the pathogenesis of lung lesions, and provide insights into host- bacteria interactions. Virulence factors should aid pathogens to quickly adapt to shifts in the environment, and improve the adhesion and invasion of pathogens to host cells^18^. It also helps pathogens to escape from the host’s innate and adaptive immune^29^. For example, *Mycobacterium tuberculosis* is a pathogen in human respiratory tract diseases. The VFs produced by *M. tuberculosis* are involved in the interaction with the host macrophages, which helps the *M. tuberculosis* to escape the host immune responses^30^. Here, 229 VFGs were identified in 285 *M. hyopneumoniae* genomes. Given *M. hyopneumoniae* as a pathogen of swine respiratory tract diseases, we inferred that VFGs should be involved in respiratory tract diseases, e.g., lung infections and lesions.

The construction of MAGs significantly increased the number of microbial genomes. This could be used to obtain strain-specific genome information^31^ and should facilitate the pan-genome analysis. *M. hyopneumoniae* is a respiratory pathogen that can cause endemic pneumonia in pigs and leads to huge economic losses for pig farms^5^. Only 23 *M. hyopneumoniae* genomes are available in the NCBI RefSeq database. A previous study revealed genetic variations in *M. hyopneumoniae* based on 16 selected genes^32^, but whole-genome surveys have not been carried out. Here, for the first time, we reported the genetic diversity of *M. hyopneumoniae* strains in porcine respiratory microbiome through an in-depth survey at both inter-individual and population-level. The number of pan-genes was more than four times than the number of genes in the present reference genome. Moreover, a large number of pan- genes could not match to COG categories. This was consistent with the reports in which the functions of up to 30% genes are still unknown in the present reference genome^16^. These results exhibited an extremely high degree of individual variation and genetic diversity in *M. hyopneumoniae* strains. Furthermore, *M. hyopneumoniae* genomes downloaded from the reference database were genetically similar and clustered together. However, *M. hyopneumoniae* MAGs reconstructed in this study were widely distributed in both MDS analysis and the phylogenetic tree (**Fig. 3c and 3d**). This further suggested the high genetic diversity of *M. hyopneumoniae* strains obtained in this study due to the large sample size and a wide sample source. We provided a large-scale of new source of *M. hyopneumoniae* genomes that would facilitate the investigation of the roles of *M. hyopneumoniae* in lower respiratory diseases.

The α-diversity of lung microbiome decreased with the severity of lung lesions. High microbial diversity had a favorable effect on maintaining homeostasis of the micro-ecosystem, whereas low diversity of the microbial composition was associated with clinical features of pneumonia in the human lung microbiome^24^. We previously found that *Mycoplasma* was associated with lung lesions by 16S rRNA sequencing analysis^26^. Here the relative abundance of *M. hyopneumoniae* increased with the severity of the lung lesions. It has been shown that *M. pneumoniae* can decrease the abundances of other bacteria by direct competition in the human respiratory microbiome^9^. Consistently, *M. hyopneumoniae* was negatively correlated with eight bacterial species in the SVLL group (**Fig. 6d**). Therefore, we suggested that the high abundance of *M. hyopneumoniae* decreased the α-diversity of the lung microbiome through decreasing the abundance of other bacteria. Several *M. hyopneumoniae* MAGs was significantly associated with lung lesions in both F_7_ pigs and the Berkshire × Licha cross lines. This result suggested the significant role of *M. hyopneumoniae* strains in lung lesions. *M. hyopneumoniae* has been known as a pathogen that attaches to the cilia and surface of epithelium in colonizing the host, resulting in damage and loss of cilia and even epithelial cell death^5^. A comparative analysis of adhesin proteoforms endoproteolytic processing between pathogenic and non-pathogenic *M. hyopnemoniae* strains demonstrated that adhesin proteins were associated with pathogenicity of *M. hyopnemoniae*^33^. Among eight VF types harbored by *M. hyopneumoniae* and enriched in lung lesion pigs (**Supplementary Fig. 9b**), seven were related to adhesion function. Furthermore, all *M. hyopneumoniae* MAGs enriched in lung lesion pigs carried at least five of these seven VF types. Thus, we speculated that adhesion-related VFs might promote the adhesion and overgrowth of pathogenic *M. hyopneumoniae* strains in host lung, and finally resulted in lung lesions. In conclusion, we constructed a non-redundant gene catalog that contained an unprecedented number of 10,337,194 genes (PRGC90) and recovered 397 MAGs from the swine lower respiratory tract microbiome. Using these catalogs, we found the heterogeneity among individuals in the microbial composition and potential functional capacities of the swine lower respiratory tract microbiome. We analyzed the pan-genome and phylogenetic structure of *M. hyopneumoniae*, and found that high abundances of *M. hyopneumoniae* strains and the adhesion-related VFs carried in their genomes were significantly related to swine lung lesions. The results from this study provide important resources for the researches on the swine lower respiratory tract microbiome and would benefit the diagnosis and treatment of lung diseases.

## Methods

### Experimental animals and sample collection

All subjects used in this study included 618 pigs from a mosaic population, 28 individuals from a Berkshire × Licha cross line, 11 Tibetan pigs, nine Erhualian pigs, and nine wild boars (**Supplementary Table 1**). The mosaic population was constructed by intercrossing four Western breeds (Duroc, Landrace, Large White, and Pietrain) and four Chinese breeds (Bamaxiang, Erhualian, Laiwu, and Tibetan). All pigs were raised under uniform indoor conditions at the experimental farm in Nanchang, Jiangxi Province. A total of 688 samples were collected from 618 F_7_ pigs of this mosaic population at the age of 240 days with sterile phosphate-buffered saline (PBS) after slaughter, including 613 bronchoalveolar lavage (BAL) samples and 74 trachea-lavage samples. As a part of the upper respiratory tract, only one lavage sample was obtained from esophagus in one of 618 F_7_ pigs, to increase the comprehensiveness and representativeness of the microbial gene and MAG catalogs of the swine respiratory tract microbiome, it was also included in the construction of the catalogs, but not used for other analyses. Twenty-eight pigs from a commercial Berkshire × Licha line raised in a commercial farm in Dingnan, Jiangxi Province were slaughtered, and 28 BAL samples were obtained. BAL samples were also collected from 11 Tibetan pigs which were initially raised on semi-free conditions in Ganzi, Sichuan Province, and then transferred to a Nanchang farm and were further raised one to three months. BAL samples from nine Erhualian pigs were also included in this study. These Erhualian pigs were raised in a farm located in Changzhou, Jiangsu province. We collected nine BAL samples from wild boars that were captured from the mountains in Jiangxi Province. All pigs received no antibiotic treatment for two months prior to sample collection. Several procedures had been taken to control the possible contaminations as far as possible: 1) sterile phosphate-buffered saline (PBS) and 50-ml sterile centrifuge tubes were separately prepared for each sample, and aseptically and closely stored. 2) After removed from the chest cavity, the lungs were placed in a clean place immediately for lavage. 3) All operators wore masks and disposable medical gloves. About 45-50ml PBS lavage from the lung was collected for each respiratory samples within 30 min after slaughter. PBS from the same batch and experienced the same process of sampling but not used for lavage was chosen as blank control. All samples were taken back to the laboratory on ice within 1 hour. The PBS lavage was centrifuged at 4000 g for 30 minutes at 4°C in a laminar flow cabinet (Esco Lifesciences, Singapore). The lavage pellet was subsequently transferred into a 2-mL sterile tube and stored at –80°C until DNA extraction for subsequent analysis.

### Phenotype measurement

Porcine respiratory diseases are considered the most important health problems in pig production. However, diagnosis of respiratory diseases in pigs is often challenging and should rely on the combination of the evaluation of lung lesions on abattoir inspection. The degree of lung lesions can be treated as an important trait to assess lung diseases in finishing pigs at slaughter^34^. Here we adopted a scoring system to evaluate lung lesions of experimental pigs at the slaughter house. In brief, the anterior and posterior views of each lung were photographed using a digital camera. Subsequently, the degree of lesions to each lung was scored using these photographs according to the methods and scoring criteria described in detailed in our previous study^35^: (I) The anterior of the lung was divided into six sections, the left and right apical lobe, left and right cardiac lobe, and left and right diaphragmatic lobe. The posterior of the lung was divided into seven sections, the left and right apical lobe, left and right cardiac lobe, left and right diaphragmatic lobe, and intermediate lobe (**Fig. 5**). For each section, the score was assigned from zero to five, corresponding to the proportion of lesion area: 0%, 0–20%, 20–50%, 50–75%, 75–90% and more than 90%. (II) Different sections were endowed with different weights according to the area of each section. The weight for left and right apical lobe, left and right cardiac lobe, left and right diaphragmatic lobe, and accessory lobe were 20% (10% for each of left and right sections), 20%, 50%, and 10%, respectively. Finally, the summed score of the anterior lung × 50% + the summed score of six sections of the posterior lung corresponding to the anterior lung × 50% + accessory lobe score × 10% was treated as the final score of lung lesions for each experimental pig. Pigs with a final score ranging from 0 to 0.75 were classified into the healthy lung group (HL), from 0.75 to 1.50 into slight lung lesion group (SLL), from 1.5 and 3.00 into moderate lung lesion group (MLL), and > 3 into the severe lung lesion group (SVLL). A total of six experienced panelists independently participated in the scoring. And then, three out of six scoring results with the highest pairwise correlation coefficient (> 0.8) were used. The mean of the scores from these three panelists was treated as the lung lesion score of each animal. Re-scoring had to be conducted if there were no three scoring results that met the requirements. A total of 675 experimental pigs were well phenotyped.

### Microbial DNA extraction and metagenomic sequencing

Microbial DNA was extracted from lavage pellets using QIAamp DNA Stool Mini Kit (Qiagen, Hilden, Germany) according to the manufacturer’s instructions. The process of DNA extraction was carried out in a laminar flow cabinet (Esco Lifesciences, Singapore). The concentration and the quality of DNA samples were measured with a NanoDrop-1000 and by 0.8% agarose gel electrophoresis. Metagenomic sequencing libraries were constructed using the VAHTS Universal DNA Library Prep Kit (Vazyme, China) following the manufacturer’s instructions. In brief, DNA samples were randomly fragmented by sonication to a size of 300 bp. And then, DNA fragments were end-polished and A-tailed at the 3′ end. Adaptors were ligated to the end of fragments for PCR amplification. After purification, all libraries were sequenced with the 150 bp paired-end strategy on a DNBSEQ-T7 platform (BGI, China). To look into the possible contamination introduced during sample collection and sequencing, we collected six PBS samples from the same batch of PBS solution for lavage as blank controls and six samples of mixed regents that were used for library construction and sequencing as sequencing background control samples. High throughput sequencing was performed for all these 12 samples. Because DNA concentrations obtained from blank control samples were far from enough for library construction, these DNA samples were condensed to obtain sufficient material for library construction and sequencing.

### Metagenomic sequence assembly

The adapters and low-quality sequences were filtered from raw data using fastp (v0.20.1, --cut_by_quality3 -W 4 -M 20 -n 5 -c -l 50 -w 3)^36^. Host genomic DNA sequence contaminations were subsequently removed from sequencing data by mapping reads to the pig reference genome (Sscrofa11.1) using Bowtie2 (v2.3.5.1)^37^ with default parameters. The co-assembly of 745 metagenomic sequencing data was employed by MEGAHIT (v1.2.9, --min-contig-len 500)^38^. Single-sample assemblies were also performed for 745 metagenomic sequencing data using MEGAHIT^38^ with the same parameters for co-assembly.

### Construction of the microbial gene catalog

Assembled contigs from both co-assembly and single-sample assembly were used for gene prediction by Prodigal (v2.6.3)^39^ with the parameter “-p meta”. Those genes with length ≥ 100bp were retained for further analyses, and genes that had lengths of < 100bp but having a complete open reading frame (with both start and stop codon) were also retained. All genes were clustered at the protein level using CD-HIT (v4.8.1)^40^ following UniRef guidelines at 100% (PRGC100), 90% (PRGC90), and 50% (PRGC50) of amino acid sequence identity. Those genes belonging to eukaryotes (except fungi and protists) were excluded from the gene catalog by aligning all genes to the Uniprot TrEMBL database (https://www.uniprot.org/statistics/TrEMBL).

### Taxonomic and functional annotation of genes

The taxonomic assignment for genes in the catalog was performed based on their amino acid sequences of the proteins using DIAMOND (v2.0.12.150)^41^ against the Uniprot TrEMBL protein database with the threshold of e-value ≤ 1 × 10 ^-5^. Proteins that were not aligned to the database were defined as unknown proteins. For genes with multiple records in the output of DIAMOND^41^, the taxonomic classifications were determined based on the lowest common ancestor algorithms by BASTA (v1.3.2.3)^42^ with the options of an alignment length > 25, and shared by at least 60% of hits. Similarly, proteins that were not assigned to any taxon were defined as unknown taxa. The eggNOG orthologs and COG functional categories were annotated by aligning genes to the eggNOG database (v5.0) using eggNOG-mapper (v2.6.1)^43^. The annotations of KEGG orthologs were obtained with KOBAS (v3.0.3)^44^ software (-t blastout:tab, -s ko). Genes in the catalog were aligned to against the dbCAN2 database (HMMdb V10) to obtain Carbohydrate-active enzymes (CAZymes) annotation using HMMER (v3.1b2)^45^. The protein amino acid sequences of genes were aligned to the Virulence Factor Database (VFDB)^18^ to obtain virulence factor annotation using BLAST (v2.12.0)^46^. Host bacteria of virulence factor genes (VFGs) were determined by aligning gene sequences to both VFDB and Uniprot TrEMBL protein database. If a gene sequence was simultaneously mapped to a VFG and a microbial taxon, we considered that microbial taxon as the host bacterium of this VFG. The VF types refer to a class of virulence factors encoded by the same kind of VFGs.

The procedure for functional annotation of MAGs was as same as that for gene annotation. For all functional alignments of protein sequences, the annotated hit(s) with the highest scores were selected for the subsequent analyses^47^.

### Metagenomic binning

Three binning software programs, MetaBAT2 (v2.15)^48^, Maxbin2 (v2.2.7)^49^, and CONCOCT (v1.0.0)^50^ were used for binning assembled contigs from single sample. Bin sets obtained from three programs were combined and refined with the Binning_refiner module of MetaWRAP (v1.3.2)^51^. The quality of bins was evaluated using CheckM (v1.0.18)^52^. The metagenome-assembled genomes (MAGs) with ≥ 50% of completeness and ≤ 10% contamination were retained for subsequent analyses. To improve the quality of MAGs, metagenomic sequencing reads mapped to MAGs were reassembled with metaSPAdes (v3.13.0)^53^ using the Reassemble_bins module of MetaWRAP^51^.To retrieve more MAGs, and considering the low sequencing depth after removing host genomic DNA contamination, a co-binning strategy was used in this study. MetaBAT2 (v2.15), Maxbin2 (v2.2.7), and VAMB (v3.0.2)^54^ were used for co-binning with the procedures applied to single-sample binning. All MAGs were dereplicated using dRep (v3.2.2)^55^, and clustered at the threshold of 99% average nucleotide identity (ANI) (strains) and 95% ANI (species-level genome bins, SGBs).Taxonomic annotation of MAGs was performed by GTDB-Tk (v1.7.0) ^15^ based on the Genome Taxonomy Database (GTDB, https://gtdb.ecogenomic.org/). These SGBs containing at least one reference genome in the GTDB were considered to be known SGBs, and SGBs that could not match to any reference genome in the GTDB were defined as unknown SGBs (uSGBs)^56^. The genome annotations of MAGs were performed using Prokka (v1.13)^57^ with default parameters, except for the MAGs classified into *Mycoplasma* and *Ureaplasma* that were annotated with the option “-- gcode 4”, because in most of the bacteria from *Mycoplasma* and *Ureaplasma*, the general UGA stop codon encodes tryptophan in the translation using genetic code 4.

### Estimation of the abundances of genes, taxa, functional terms, and MAGs

Clean reads of each sample were mapped to the gene catalog using BWA MEM2 (v2.2.1)^58^. Samtools (v1.7)^59^ was used for converting samfiles to bamfiles and for sorting. The counts of successfully assigned reads were calculated by FeatureCounts (v2.0.1)^60^. The relative abundance of each gene in each sample was calculated by the following formula:

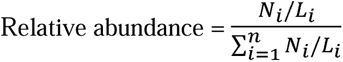

where for gene *i, N_i_* is the number of reads mapped to gene *i*, and *L_i_* represents the sequence length of gene *i*, n is the number of genes per sample. The relative abundance of taxa and functional categories were calculated by summing the relative abundances of genes which assigned to each category. The abundance of MAGs in each sample was calculated by the Quant_bins module in metaWRAP (v1.3.2)^51^.

## Phylogenetic Analysis

The Phylogenetic tree of all MAGs was constructed by PhyloPhlAn (v3.0.60)^61^ based on 400 universal marker proteins with the parameters: “--diversity low -- min_num_marker 50”. The other softwares and the options involved in this process are listed below:

- Diamond (v0.9.24.125)^62^ modules used in this study included ‘‘Blastx’’ for the nucleotide-based mapping, and “Blastp’’ for the amino acid based mapping. The parameters ‘‘--outfmt 6 --more-sensitive --id 50 --max-hsps 35 -k 0’’ were used in both modules;
- MUSCLE (v3.8.1551)^63^, with ‘‘–-quiet -maxiters 2’’ options;
- trimAl (v1.4.rev15)^64^, with ‘‘-gappyout’’ option;
- IQ-TREE (v2.2.0-beta)^65^, with ‘‘-quiet -nt AUTO -m LG’’ options;
- RAxML (v8.2.12)^66^, with ‘‘-p 1989 -m GTRCAT’’ options.

The phylogenetic tree for 285 *M. hyopneumonia* was built with PhyloPhlAn using 401 core genes (present in more than 95% of the genomes) that were determined by Roary (v3.12.0, -e -z -i 95 -cd 95 -t 4)^67^. The parameters were set as follows: “--diversity low --trim greedy --remove_fragmentary_entries”. The software programs and the parameters involved in this process are listed follows:

- Blastn (v2.9.0+)^46^, with ‘‘-outfmt 6 -max_target_seqs 1000000 -perc_identity 75’’ options;
- MAFFT (v7.490)^68^, with “--anysymbol --auto” options;
- FastTree (v2.1.10)^69^, with “-quiet -pseudo -spr 4 -mlacc 2 -slownni -fastest -no2nd

- mlnni 4 -gtr -nt” options;
- RAxM and trimAl were set with the same options as mentioned above.

The phylogenetic tree for all MAGs was visualized using iTOL (v6.5.2)^70^, and the phylogenetic tree for 285 *M. hyopneumoniae* and the taxonomic tree for lung core taxa were visualized by GraPhlAn (v1.1.3)^71^.

### Pan-genome analysis of *M. hyopneumoniae*

We downloaded 23 genomes labeled as *M. hyopneumoniae* from the NCBI RefSeq repository. MAGs from metagenomic single-sample binning analysis annotated as *M. hyopneumoniae* were included in the pan-genome analysis. These genomes were quality controlled by a series of strategies. Firstly, quality control was performed for all *M. hyopneumoniae* MAGs. The MAGs meeting the following standards were retained for further analysis: 0.7 Mbps < genomes size < 1.1 Mbps (because the size of 23 public *M. hyopneumoniae* genomes downloaded from the NCBI RefSeq repository ranged from 0.8 - 1 Mbps), containing < 500 contigs, >90% completeness and < 5% contamination. Finally, a total of 285 *M. hyopneumoniae* genomes (23 reference genomes and 262 MAGs) passed the quality control criteria and were used for the pan-genome analysis. Prokka (v1.13, --gcode 4)^57^ was then used to annotate these genomes. The annotated genomes were processed with Roary pipeline^67^. Those genes present in > 95% of the genomes and having a minimum 95% identity were defined as core genes and were used for phylogenetic analyses. To define sub-species of *M. hyopneumoniae*, pairwise ANI values between genomes were calculated using FastANI (v1.32)^72^. To determine the optimal number of sub-species, the requirement of prediction strength (prediction.strength function in *fpc* package) > 0.8 was applied^73^. The partitioning around medoids (PAM) clustering algorithm was used to cluster genomes to distinct sub-species using the *cluster* package in R. The sub-clade was defined using *rhierbaps*^74, 75^ in R (v4.1.2) package based on the concatenated nucleotide core gene alignment that was produced by PhyloPhlAn. Classical Multidimensional Scaling (MDS) analysis was conducted based on the pairwise ANI distance matrix using the cmdscale function in R.

### Statistical analysis

Accumulation curve of the number of predicted genes in the gene catalog were bootstrapped 10 times at each sampling interval. Asymptotic regressions were carried out using the SSasymp and nls functions in the R *stats* package. Linear discriminant analysis effect size (LEfSe) analysis (v1.1.01)^76^ was used to identify MAGs, virulence factors, and KEGG pathways showing different enrichments among pig groups with different severity of lung lesions in F_7_ pigs, and between healthy and lung lesion pigs in the Berkshire × Licha cross line at the threshold of |LDA| >2. The α-diversity indices including observed species and the Shannon index, and the β-diversity metric of the lung microbial composition based on Bray-Curtis distance were calculated with the R *vegan* package^77^. Principal coordinate analysis (PCoA) was also performed with the R *vegan* packages using Bray–Curtis dissimilarities. The comparison of functional capacities between two *M. hyopneumoniae* sub-species, and the comparisons of the α- diversity and lung microbial composition among four pig groups with different severity of lung lesions were carried out using Wilcoxon tests. The co-occurrence network of 27 bacterial species in each pig group was constructed based on the Spearman rank correlations in relative abundance among bacterial species. The *P* values were calculated based on 999 permutations. Only the interactions between bacteria species with correlation coefficient > 0.3 and *P* < 0.05 were present in the co-occurrence network. The co-occurrence networks were visualized and annotated in Cytoscape (v3.9.1)^78^.

## Data availability

All metagenomic sequencing data are available in the GSA database (https://ngdc.cncb.ac.cn/gsa/browse/CRA007668) under accession number CRA007668. All MAGs and microbial genes in the gene catalog were also deposited in the GSA database (https://ngdc.cncb.ac.cn/bioproject/browse/PRJCA010893) with accession numbers GWHBPMO00000000∼ GWHBQBU00000000 and OMIX002571, respectively.

## Code availability

The source data are available at GitHub (https://github.com/Jingquan-Li/PRGC).

## Acknowledgments

We thank the colleagues in the State Key Laboratory of Pig Genetic Improvement and Production Technology, for their technical supports in managements of experimental pigs and the sample collections. This work was supported by China Agriculture Research System (No. CARS-35).

## Author contributions

L.H. conceived and supervised the study, interpreted the results and revised the manuscript; C.C. co-supervised the study, interpreted the results and revised the manuscript; H.A. conceived the study and collected the samples; J.L. collected the samples, phenotyped lung lesions, performed metagenomic analysis and statistical analysis, visualized the data and wrote the manuscript; F.H. collected the samples, phenotyped lung lesions and wrote the manuscript; Y.Z. performed metagenomic analysis and statistical analysis; T.H., X.T., M.Z., J.C., Z.Z., J.L., F.H., H.D., Z.L., M.Z., and Y.X phenotyped lung lesions and collected the samples. All authors read and approved the final manuscript.

## Materials & Correspondence

Correspondence and requests for materials should be addressed to L.H.

## Supplemental information

Supplementary Figures and the Supplementary Tables with small size (within one page) including Supplementary Table 2, Supplementary Table 4, Supplementary Table 7, and Supplementary Table 10 were provided as a single combined PDF document. Supplementary Tables including Supplementary Tables 1, 3, 5-6, and 8-9, whose sizes exceed more than one page were provided as an Excel file type.

